# Inhibitory networks orchestrate the self-organization of computational function in cortical microcircuit motifs through STDP

**DOI:** 10.1101/228759

**Authors:** Stefan Häusler, Wolfgang Maass

## Abstract

Interneurons have diverse morphological and physiological characteristics that potentially contribute to the emergence of powerful computational properties of cortical networks. We investigate the functional role of inhibitory subnetworks in the arguably most common network motif of cortical microcircuits: ensembles of pyramidal cells (PCs) with lateral inhibition, commonly referred to as Winner-Take-All networks. Recent theoretical work has shown that spike-timing-dependent plasticity installs in this network motif an important and ubiquitously useful self-organization process: The emergence of sparse codes and Bayesian inference for repeatedly occurring high-dimensional input patterns. However, this link has so far only been established for strongly simplified models with a symbolic implementation of lateral inhibition, rather than through the interaction of PCs with known types of interneurons. We close this gap in this article, and show that the interaction of PCs with two types of inhibitory networks, that reflect salient properties of somatic-targeting neurons (e.g. basket cells) and dendritic-targeting neurons (e.g. Martinotti cells), provides a good approximation to the theoretically optimal lateral inhibition needed for the self-organization of these network motifs. We provide a step towards unraveling the functional roles of interacting networks of excitatory and inhibitory neurons from the perspective of emergent neural computation.

## 1 Introduction

The fundamental organization of cortical processing is still unknown. Increasing evidence indicates that the brain relies on a set of canonical neural networks, repeating them across brain regions and modalities to apply similar operations to different problems. Douglas and Martin (2004) proposed to view winner-take-all (WTA) networks, where pyramidal cells (PCs) inhibit each other via interneurons, as an essential canonical network motif. Several theoretical and modeling studies have demonstrated that powerful computational functions can emerge in these network motifs through synaptic plasticity. In particular, competitive Hebbian learning has the potential to induce the self-organization of probabilistic models and sparse coding of complex inputs to WTA networks (Rumelhart and Zipser, 1988; Nowlan, 1990, 1991; Neal and Hinton, 1998; Sato, 1999; Keck et al., 2012). Moreover, it has recently been shown (Habenschuss et al., 2012; Nessler et al., 2013) that spike-timing dependent plasticity (STDP) is able to approximate the arguably most powerful known learning principle for learning a probabilistic model for the spiking activity of upstream neurons of WTA networks: Expectation Maximization.

Although this work points to a powerful role of inhibitory neurons for the emergence of generic computational properties of cortical microcircuits, it had bypassed the question of their specific role in this context. The interaction of excitatory and inhibitory neurons was replaced by a symbolic and theoretically ideal term for lateral inhibition. In contrast, interneurons have diverse morphological and physiological characteristics (Markram et al., 2004; Ascoli et al., 2008) that potentially result in distinct functional roles. In particular, PCs receive inhibitory inputs from two main types of inhibitory neurons, i.e. dendritic and somatic-targeting interneurons. About 50% of all inhibitory interneurons are basket cells, which specialize to target the somata and proximal dendrites of PCs. Their primary function is to control the gain of the integrated synaptic response. The largest class of dendritic-targeting interneurons are somatostatin expressing Martinotti cells. These interneurons specialize to target mainly distal dendrites and tuft dendrites of PCs. Interneurons that preferentially target the dendritic domain are positioned to influence dendritic processing and synaptic plasticity.

Recent theoretical work (Körding and König, 2000, 2001) proposes specific functional roles for each of these inhibitory neuron types: Somatic-targeting interneurons define the activity of ensembles of PCs, whereas dendritic-targeting interneurons gate synaptic plasticity at excitatory synapses onto PCs. We argue that this approach solves an intrinsic problem of previous models that implement probabilistic inference in WTA networks: The requirement of a normalized firing rate of PCs that can neither be achieved by feedforward inhibition nor by delayed feedback inhibition. In particular, we show in simulation that the ideal WTA network model proposed in (Habenschuss et al., 2012), that is based on an ideal symbolic inhibition and normalized firing rates of PCs, can be implemented with spiking interneurons and unnormalized firing rates if an additional gating mechanism for synaptic plasticity is implemented with dendritic-targeting interneurons.

We test the performance of WTA networks on a realistic self-organization task: Sparse coding of two classes of firing rate patterns that occur repeatedly in the input stream, where the patterns in one class occur only half as often as the patterns in the other class. This task provides a hard challenge for any self-organization approach, since in most approaches more neurons respond to the more frequently occurring patterns, and possibly none to the more rarely occurring patterns. We show that a combination of both types of interneurons, i.e. basket cells and Martinotti cells, achieves the best approximation to the theoretically optimal self-organization of the WTA network after a low number of one hundred pattern repetitions.

We verify that the resulting WTA networks are consistent with experimental data on the weak but non-negligible correlation of PCs in superficial layers of sensory cortex. We demonstrate that the resulting WTA networks are tolerant to a wide range of synaptic delays and connection probabilities, and are therefore consistent with experimental data on the fine structure of various cortical microcircuits. Altogether, we provide a contribution towards closing the gap between biological data on diverse types of interneurons in cortical networks and their putative computational function.

## 2 Materials and Methods

### 2.1 WTA network

This section briefly summarizes the WTA model in (Habenschuss et al., 2012) (see supp. material for details). The WTA network model consists of a population of PCs z_1_, …, z*_K_* that inhibit each other by means of lateral inhibitory connections (Fig. 1). The *k*th PC in the WTA network spikes at time *t* in response to synaptic input from upstream neurons y_1_, …, y*_N_* with a firing rate

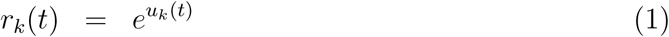

that depends exponentially on its membrane potential *u_k_ (t)*. Such an exponential dependence of the firing rate on the membrane potential is consistent with experimental data (Jolivet et al., 2006). The membrane potential *u_k_(t)* of neuron *z_k_* is given by

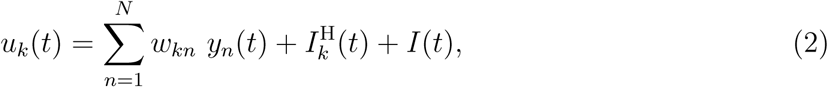

where *w_kn_* is the weight of the synaptic connection between upstream neuron *y_n_* and PC z*_k_*. 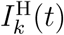 and *I(t)* denote two additional inhibitory potentials (see below) and *y_n_* denotes the excitatory post-synaptic potential (EPSP) triggered by upstream neuron *y_n_*. The spikes of upstream neuron *y_n_* are modeled as a Poisson process. The total firing rate of all PCs in the WTA network is given by

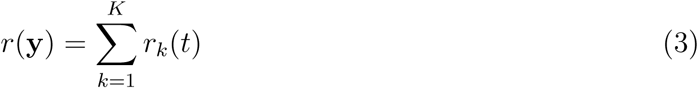

and depends on the EPSPs **y**.

**Figure 1:**
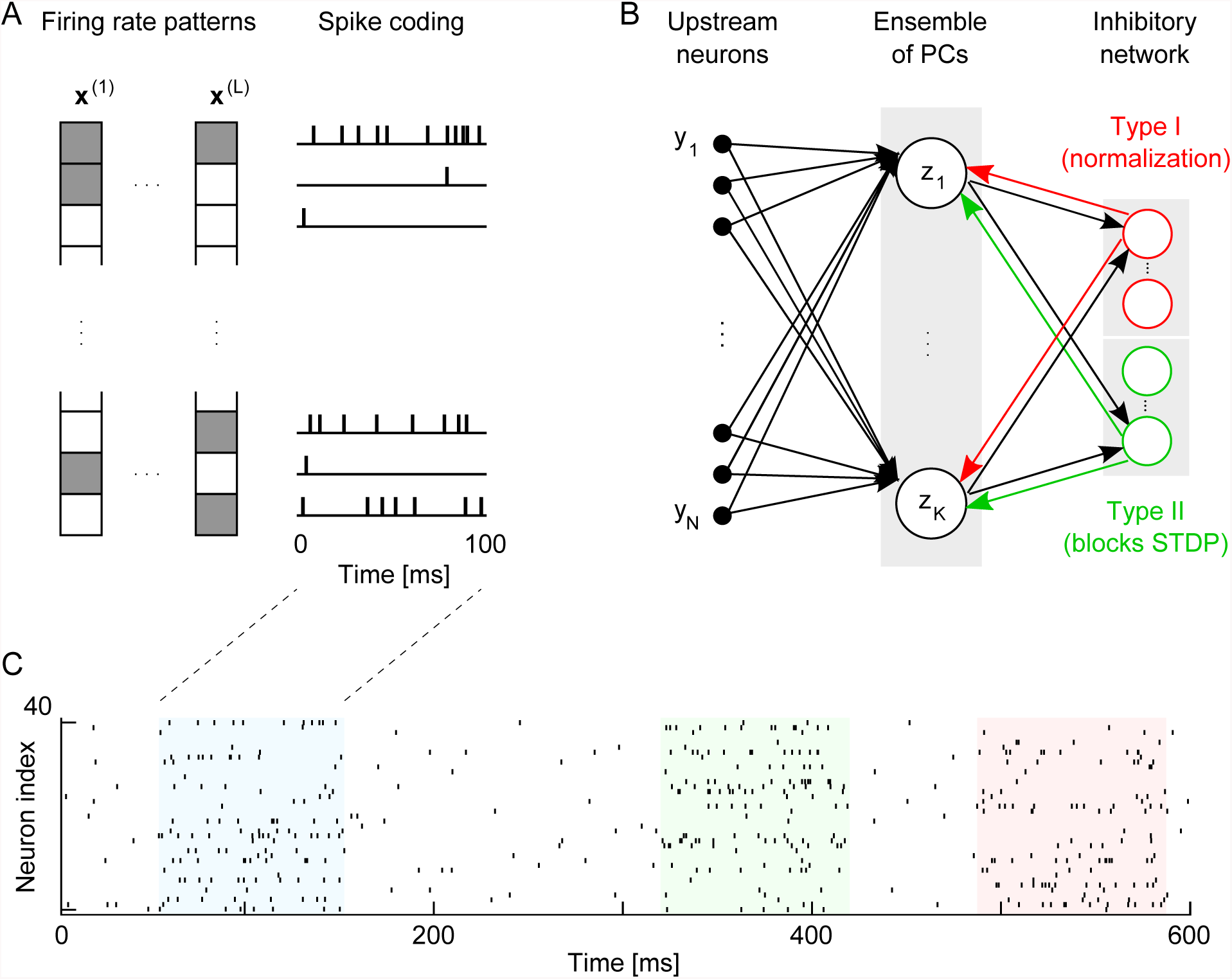
WTA network architecture that induces probabilistic inference. **A:** Firing rate patterns **x**^*(l)*^ with *l = 1,…, L.* Each element of a pattern defines for each upstream neuron y_1_, …, yN the firing rate of a homogeneous Poisson spike train. White and gray elements denote firing rates of 10 Hz and 100 Hz, respectively. Poisson spike trains with corresponding firing rates and a duration of 100 ms are injected into the PCs z_1_, …, z*_k_*. **B:** Modification of the WTA network proposed in (Nessler et al., 2013; Habenschuss et al., 2012). The standard WTA network (shown in black) is extended by two inhibitory networks (marked in green and red). The first inhibitory network consists of a population of inhibitory neurons that provides inhibition of type I (marked in red). The second population of inhibitory neurons provides inhibition of type II (marked in green). All PCs in the WTA network receive the same inhibition of type I and II. **C:** PCs in the WTA network receive continuous spike input that is either generated from the firing rate patterns shown in A or from a background firing rate of 2 Hz. Colored areas indicate the presentation time intervals of firing rate patterns. Only during these time intervals the weights of excitatory synapses onto PCs are updated through STDP. We conjecture that this update is gated by neuromodulators like Acetylcholine.

The inhibitory potential 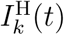 implements a mechanism that is strongly reminiscent of homeostatic intrinsic plasticity in cortex (Desai et al., 1999; Watt and Desai, 2010).It normalizes the firing rate of each PC averaged over time (see supp. material for details). The inhibitory potential *I(t)* implements global lateral inhibition that is modeled in (Habenschuss et al., 2012; Nessler et al., 2013) as an ideal and symbolic term

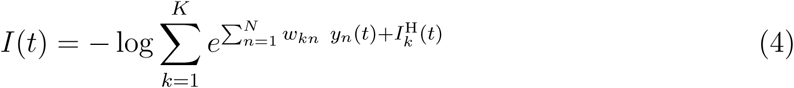

This term instantaneously normalizes the total firing rates of all PCs in the WTA network. We approximate this symbolic inhibition with a population of spiking inhibitory neurons (see Fig. 1B and Sec. 2.3), which we refer to as *inhibition of type I.*

Habenschuss et al. (2012) have shown that stochastic STDP with an expected synaptic weight change

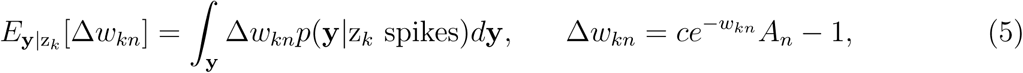

where the learning rate *c >* 0 and the contribution of the STDP learning window *A_n_* ≈ *y_n_,* in combination with precise symbolic inhibition (Equ. 4) implements a probabilistic model for the activity of upstreams neurons: The synaptic weights converge towards the fixed point

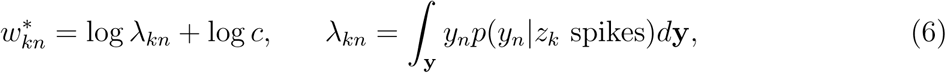

where λ*_kn_* represents the average firing rate of upstreams neuron y*_n_* at time points of post synaptic spikes of PC *z_k_*. The amplitude of the update Δ*w_kn_* depends on the value of the synaptic weight *w_kn_* before the update. According to this rule positive updates are only performed if the presynaptic neuron fired before the postsynaptic spike. It therefore respects the causality principle of long-term potentiation (LTP) that is implied in Hebbs original formulation (Hebb, 1949). At natural frequencies this update rule gives rise to a form of STDP that is very similar to plasticity curves observed in biology (Bi and Poo, 1998; Sjöström et al., 2001). We now demonstrate that this WTA network model based on normalized firing rates of PCs can be implemented with unnormalized firing rates if an additional gating mechanism for synaptic plasticity is implemented.

### 2.2 Learning with arbitrary firing rates of PCs

If the original synaptic weight changes Δ*w_kn_* are replaced by

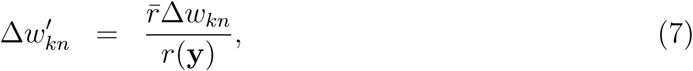

where *r̅* denotes the total firing rate averaged over time, then the expected synaptic weight changes of stochastic post-synaptically triggered STDP are invariant to the total firing rate of PCs. In particular, the fixed points of the synaptic weights 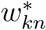 in the model of Habenschuss et al. (2012) are invariant to changes of *r(y)* as well as. The modified synaptic weight updates can be implemented by blocking a fraction of the synaptic weight updates with a probability

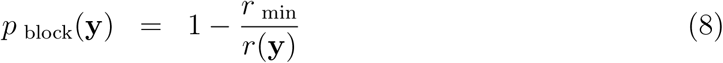

that depends on the minimum total firing rate *r_min_* = min_y_ *r*(y). Then, the expected synaptic weight changes for partially blocked STDP and unnormalized total firing rates *r*(y)

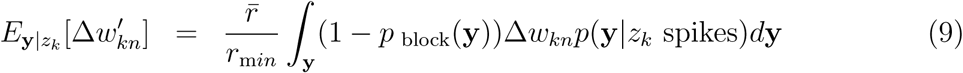

are identical to the original expected synaptic weight changes (Equ. 5) for normalized *r*(y) = *r̅*

We conjecture that the blocking of STDP is implemented by feedback inhibition that intercepts backpropagating action potentials in PCs. If backpropagating action potentials fail, then NMDA receptors may not be sufficiently depolarized to provide enough calcium influx to trigger long-term potentiation. We block STDP weight updates if at least one spike arrives during the preceding time interval of length *τ*_block_ = 6 ms from an additional inhibitory population (see Fig. 1B and Sec. 2.4), which we refer to as *inhibition of type II.* The blocking probability in Equ. 8 is obtained if the sum of the firing rates of all inhibitory neurons in this population satisfies

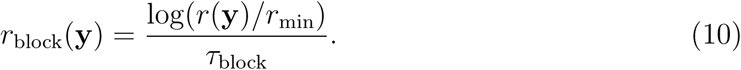

We show in Fig. 5 that that the precise shape of *r*_block_(y) is not relevant for the performance of WTA networks.

### 2.3 Inhibition of type I

Inhibition of type I is implemented with a population of spiking inhibitory neurons. The membrane potential of the *n*th inhibitory neuron is modeled as a linear sum of the EPSPs triggered by PCs, i.e.

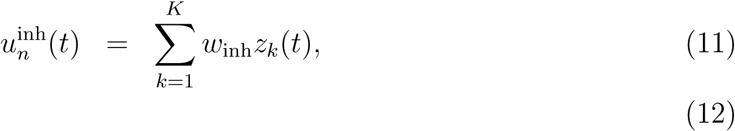

with uniform excitatory synaptic weights *w*_inh_ = 1.5 mV. The EPSPs evoked by spikes from PC z*_k_*, denoted as *z_k_*, are modeled like EPSPs in PCs evoked by upstream neurons, but with a smaller membrane decay time constant of 15 ms.

Spikes of inhibitory neurons of type I are modeled with a Poisson process with a firing rate that is proportional to 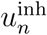, i.e.

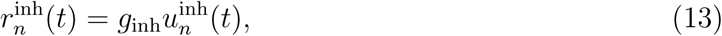

with gain *g*_inh_. The sum of the firing rates of all inhibitory neurons of type I is given by

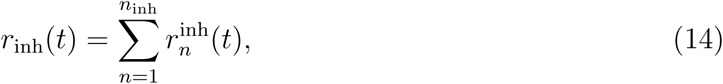

where *n*_inh_ denotes the number of inhibitory neurons of type I. The gain *g*_inh_ is set to a fixed value that results in a total firing rate *r*_inh_ that is 0, 1, 8 or 64 times *r*(y). The linear dependence of 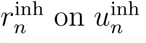 does neither constrain the number of inhibitory neurons of type I nor the size of the weights *w*_inh_, but only the product of both.

The inhibitory post-synaptic potentials 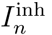 evoked by spikes from the *n*th inhibitory neuron of type I are modeled as EPSPs, but with a decay time constant of 6 ms. The total inhibition of PCs is modeled as

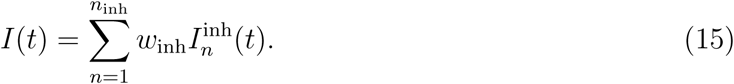

### 2.4 Inhibition of type II

Inhibition of type II is implemented with a second population of spiking inhibitory neurons. These neurons implement a mechanism that blocks STDP for excitatory synaptic connections from upstream neurons onto PCs. The membrane potential of the *n*th inhibitory neuron is modeled as a linear sum of the EPSPs triggered by PCs, i.e.

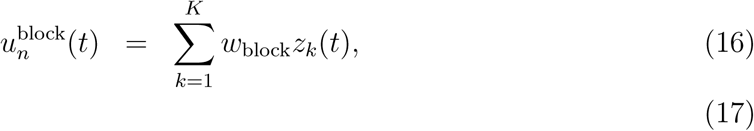

with uniform excitatory synaptic weights *W*_block_ = 1.5 mV. The EPSPs evoked by spikes from PCs are modeled as for inhibitory neurons of type I.

Spikes of inhibitory neurons of type II are modeled with a Poisson process with a firing rate that is a function *f* of its membrane potential, i.e.

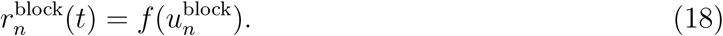

The total firing rate of all inhibitory neurons of type II is given by

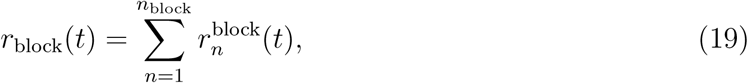

where *n*_block_ denotes the number of inhibitory neurons of type II. The function *f* is defined as

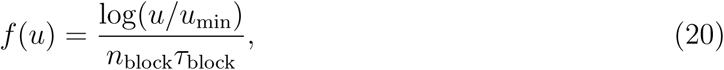

if *u > u*_min_ and set to zero otherwise. *u*_min_ denotes the mean membrane potential of an inhibitory neuron of type II if the total firing rate of PCs is *r*_min_. Simulations in Fig. 5C show that a stepwise transformation of f into a linear or an exponential function of *u* (see supp. material for details) has only a minor effect on the performance of WTA networks.

In order to disentangle the effect of both types of inhibition on the performance, the inhibitory post-synaptic potentials of synapses from the second inhibitory network onto PCs are not modeled.

## 3 Results

We investigate self-organization in the WTA network in (Habenschuss et al., 2012). In contrast to other rate-based models (Sato, 1999; Sato and Ishii, 2000; Keck et al., 2012) this model is based on spiking PCs and every spike has a clear semantic interpretation. It instantaneously assigns a hidden cause, an underlying firing rate pattern, to the spiking activity of upstream neurons. The probability of a spike depends on how well the current activity of upstream neurons agrees with the firing rate patterns encoded in the synaptic weights of PCs. These patterns are learned with STDP and optimal in the sense that they have maximum likelihood. Synaptic plasticity is carried out in combination with a homeostatic process that ensures that the number of PCs specialized to detect a particular firing rate pattern is directly proportional to the frequency of the occurrence of that pattern. The model of (Habenschuss et al., 2012) as well as related work (Sato, 1999; Sato and Ishii, 2000; Keck et al., 2012) rely on a symbolic lateral inhibition that normalizes the sum of firing rates of all PCs. We implement this inhibition with populations of spiking inhibitory neurons.

We test the performance of the WTA network on a simple self-organization task. We introduce two classes of firing rate patterns that occur repeatedly in the input stream, where the patterns in one class occur only half as often as the patterns in the other class. In total, we generate 2*K*/3 random firing rate patterns, where K denotes the number of PCs in the WTA network. Each firing rate pattern is represented by a binary vector that specifies the firing rates of 900 upstream neurons (Fig. 1A). For each vector 25% of its elements are chosen at random and set to a high firing rate of 100 Hz. The remaining elements are set to a low firing rate of 10 Hz. Each time a pattern is chosen for presentation a multi-dimensional homogeneous Poisson spike train with firing rates specified by the corresponding vector and a duration *T* = 100 ms (if not otherwise stated) is injected into the WTA network. In case of correct learning, as for the ideal model based on symbolic inhibition, rare and frequent patterns are encoded by the firing of one and two PCs. The performance of WTA networks is evaluated by estimating the percentage of correctly detected firing rate patterns after learning. A pattern is detected if there exists at least one PC that is only active during the presentation of this pattern.

### 3.1 Competition between PCs through feedback inhibition

We first investigate to what extend the ideal symbolic inhibition of the WTA network can be approximated by feedback inhibition provided by an inhibitory network (Fig. 1B). We refer to this type of inhibition as *inhibition of type I*. We assume that STDP is gated by neuromodulators (like Acetylcholine) and learning occurs only during the occurrence of firing rate patterns (colored time intervals in Fig. 1C). Therefore, we set the time intervals between firing rate patterns to zero (Supp. Fig. 1A and B).

WTA networks without feedback inhibition lack competition and all PCs detect the same firing rate pattern (Fig. 2A). The stronger the competition through feedback inhibition the better the performance (Fig. 3C). For very high firing rates of inhibitory neurons of type I (the sum of the firing rates of all inhibitory neurons averaged over time is 64 times larger then corresponding firing rate for PCs), about 60% of all firing rate patterns are detected after learning. Strong feedback inhibition causes oscillations in the total firing rate of PCs (Fig. 2C) with an approximately independent number of spikes per pattern presentation. This mechanism normalizes the total firing of PCs to some extend causing a performance improvement. The shorter the firing rate pattern duration T the better the performance (Fig. 3A and C). For short pattern durations only one oscillation of the total firing rate of PCs occurs per pattern presentation and pattern dependent differences in the oscillation frequencies have a smaller impact on the performance. In summary, the stronger the competition between PCs through inhibition of type I the better the performance.

**Figure 2:**
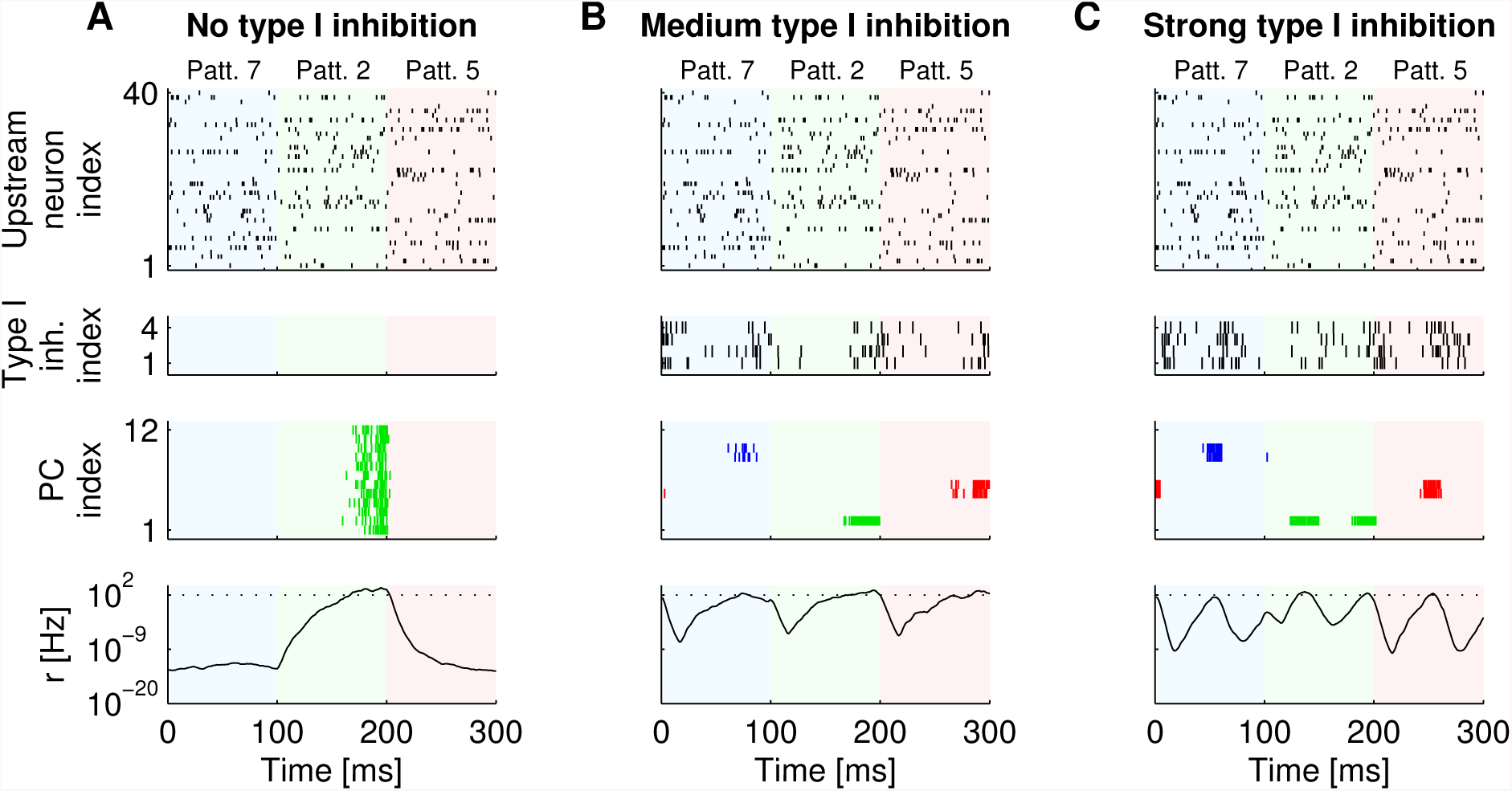
Spiking activity of WTA networks that implement inhibition of type I. **A-C:** Top and middle panels show the spike trains for 40 out of 400 upstream neurons, four inhibitory neurons providing inhibition of type I, and twelve PCs after learning, respectively. The bottom panels show the sum of the firing rates of all twelve PCs over time. The dotted line indicates a firing rate of 100 Hz. The strength of the feedback inhibition varies across panels. The average firing rate of the inhibitory population is set to 0 (A), 8 (B) or 64 (C) times the sum of the average firing rates of all PCs *r*. In the WTA network without feedback inhibition (A) all PCs detect the same firing rate pattern because of a lack of competition between PCs. Background colors indicate the indices of presented firing rate patterns. Colors of spikes indicate the index of the firing rate pattern that is detected by the corresponding PC.

**Figure 3:**
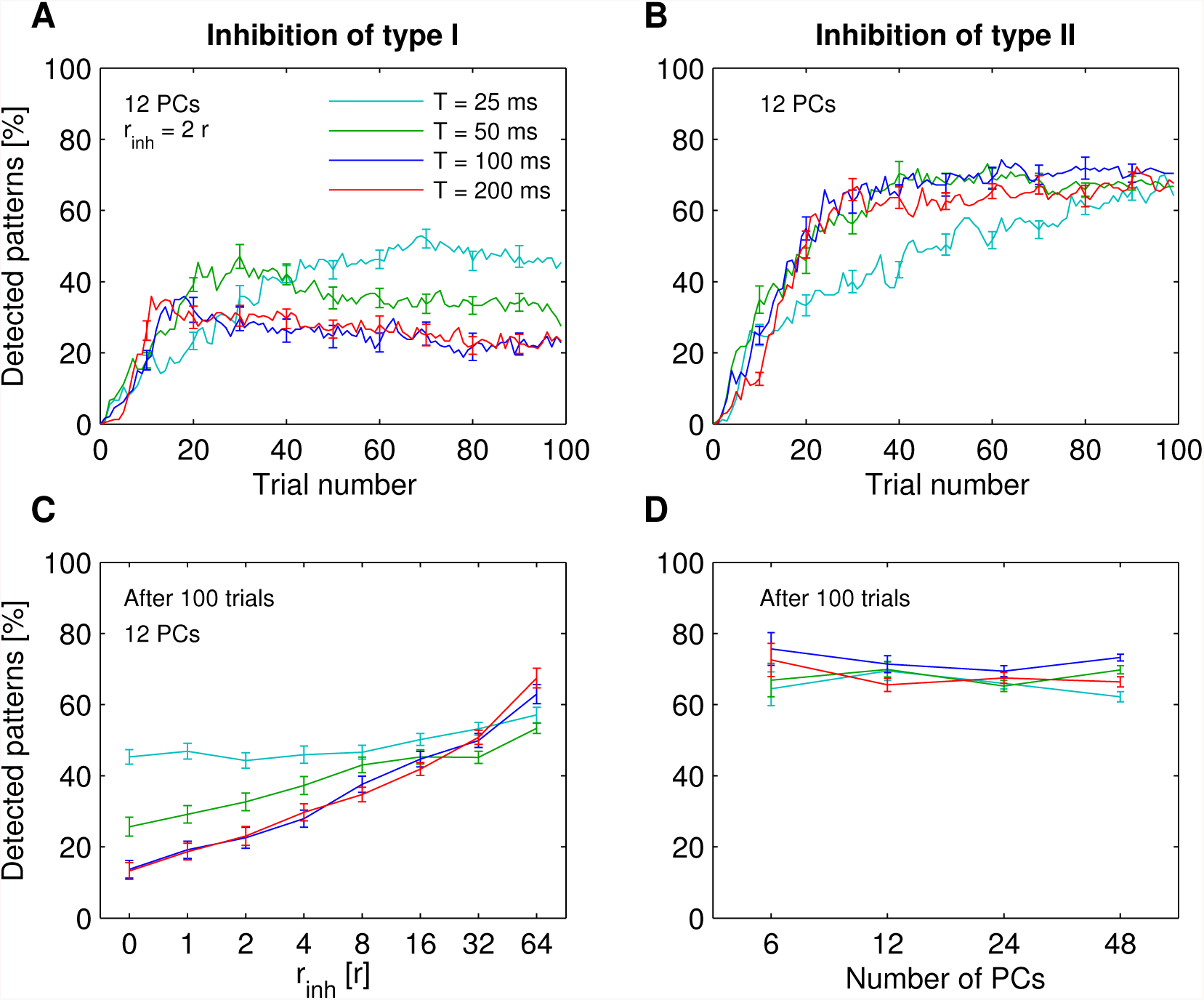
Performance of WTA networks that implement inhibition of type I or inhibition of type II. **A,B:** Percentage of detected firing rate patterns for WTA networks that implement either inhibition of type I (A) or II (B) for various pattern presentation times *T*. **C**: Percentage of detected firing rate patterns for WTA networks that implement inhibition of type I for various strengths of feedback inhibition. The average total firing rate of all neurons in the inhibitory population *r*_inh_ is set to a multiple of the average firing rate of all PCs *r*. The stronger the feedback inhibition the better the performance. **D**: Percentage of detected firing rate patterns for WTA networks that implement inhibition of type II for various pattern presentation times *T* and numbers of PCs.

### 3.2 Competition between PCs through blocking STDP

A precise normalization of the firing rates of PCs can not be achieved with spiking inhibitory neurons that provide either feedforward or feedback inhibition. Feedforward inhibition from upstream neurons can not provide precise symbolic inhibition because the strength of symbolic inhibition depends on (inaccessible) synaptic weights of excitatory synapses onto PCs that change over time (Supp. Fig. 1C and D). Feedback inhibition cannot provide precise symbolic inhibition as well. First, symbolic inhibition is provided instantaneously, whereas feedback inhibition is delayed due to synaptic transmission. Secondly, feedback inhibition depends on the total firing rate of PCs, whereas symbolic inhibition does not. Symbolic inhibition implements a constant total firing rate of PCs and depends only on the spiking activity of upstreams neurons and the weights of excitatory synapses onto PCs.

We hypothesize that excessive synaptic weight changes because of too high firing rates of PCs can be blocked by feedback inhibition. In this case, competition between PCs is not implemented by means of a normalized activity of PCs, but in form of a normalized rate of synaptic weight changes. We conjecture that STDP weight updates are blocked by means of inhibition that intercepts backpropagating action potentials in PCs. If backpropagating action potentials fail, then NMDA receptors may not be sufficiently depolarized to provide enough calcium influx to trigger long-term potentiation. This type of inhibition is presumably implemented by Martinotti cells that specialize to target mainly distal dendrites of PCs. We implement a second inhibitory network (see Fig. 1B), which we refer to as *inhibition of type II,* and model the blocking of STDP weight updates with a phenomenological mechanism: Each spike of an inhibitory neuron in this population blocks STDP weight updates of excitatory synapses onto PCs during a subsequent time interval τ_block_. In order to obtain a synaptic weight update rate that is independent of the total firing rate of PCs, r, the sum of the firing rates of all inhibitory neurons of type II has to follow

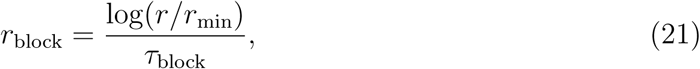

where *r*_min_ denotes the minimum value of *r*. However, we show in Sec. 3.4 that the precise shape of the firing rate function is not crucial for the performance of the WTA network.

The performance of WTA networks that implement only inhibition of type II and no inhibition of type I is shown in Fig. 3B and D. WTA networks detect more than 60% of all firing rate patterns after learning. The performance is approximately independent of the pattern duration *T* and the total number of PCs. In summary, competition between PCs through inhibition of type II results in higher performance values than competition through inhibition of type I of weak or moderate strength. However, not all firing rate patterns are detected.

### 3.3 The combination of both types of competition results in superior performance

We further investigated how well WTA networks that implement either inhibition of type I or II detect rare and frequent firing rate patterns. Fig. 4 shows that for both types of WTA networks frequent patterns are detected about twice as often as rare patterns. In particular, WTA networks that implement inhibition of type II detect about 50% of the rare patterns (without inhibition of type I, thick dark blue curve in Fig. 4B) and all frequent patterns (thick dark blue curve in Fig. 4D). Adding inhibition of type I further improves the performance for the detection of rare patterns. Nearly all rare patterns are detected for very strong feedback inhibition of type I.

**Figure 4:**
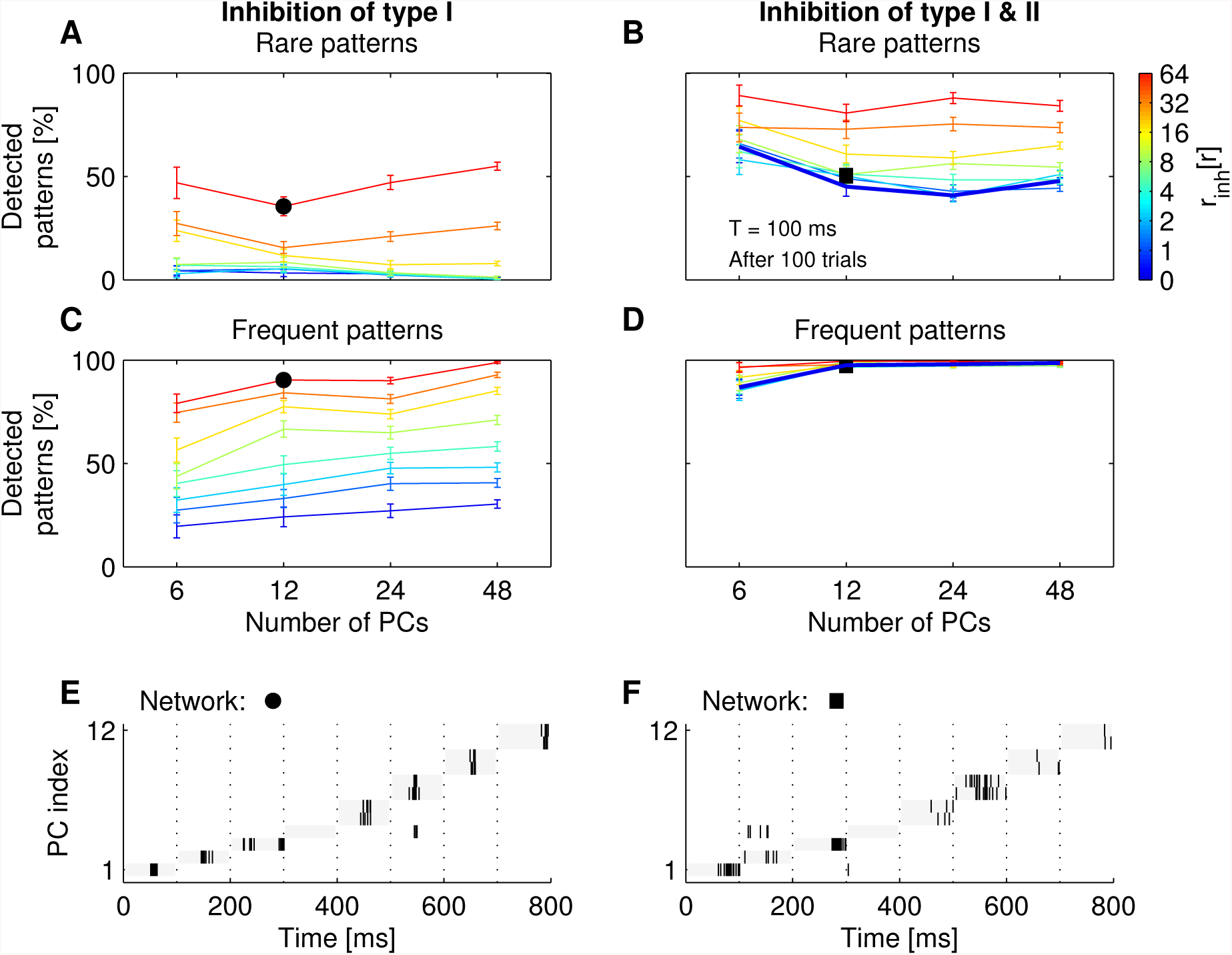
Performance of WTA networks that implement inhibition of type I and II. **A,C:** Performance of WTA networks that implement only inhibition of type I. Percentage of detected firing rate patterns that occur with a low (A) or a high frequency (C) in dependence of the strength of the feedback inhibition and the number of PCs in the WTA network. The average total firing rate of all inhibitory neurons of type I *r*_inh_ is set to a multiple of the average firing rate of all PCs *r*. **B,D:** Performance of WTA networks that implement inhibition of type I and II. Percentage of detected firing rate patterns that occur with a low (B) or a high frequency (D) in dependence of the strength of the feedback inhibition of type I and the number of PCs. The performance for WTA networks without inhibition of type I is marked with a thick dark blue curve. **E,F:** Sample spike trains for PCs of two WTA networks after learning (performance marked with a circle in A and C and a square in B and D). The duration of each firing rate pattern *T* is 100 ms (without gaps between pattern presentations). The first and the last 4 patterns are shown rarely and frequently to the WTA network during learning, respectively. Both WTA networks misclassify one rare firing rate pattern. A comparison of A and C with B and D shows that inhibition of type II improves the performance for WTA networks with no or medium inhibition of type I.

Fig. 4E and F illustrate the difference in the spiking activity of WTA networks that implement only inhibition of type I and networks that implement both types of inhibition. For the example shown, both WTA networks have similar average performance and fail to detect one rare firing rate pattern. The neural dynamics of WTA networks after learning does not change significantly, if in addition to inhibition of type I also inhibition of type II is implemented (results not shown). Finally, the temporal dynamics of excitatory synaptic weights onto PCs during learning (for WTA networks that implement both types of inhibition) resembles the one for WTA networks with symbolic inhibition (results not shown). Overall, the combination of both types of inhibition results in a performance improvement compared to WTA networks with only one of the two types of inhibition.

### 3.4 Inhibition neither has to be accurate nor precisely timed

Cortical inhibitory neurons have no (precise) logarithmic firing rate function as required for inhibitory neurons of type II. Furthermore, the synaptic transmission to and from neurons in the inhibitory populations is delayed and not instantaneous. Fig. 5 shows that both of these shortcomings have little impact on the performance of WTA networks that implement both types of inhibition.

**Figure 5:**
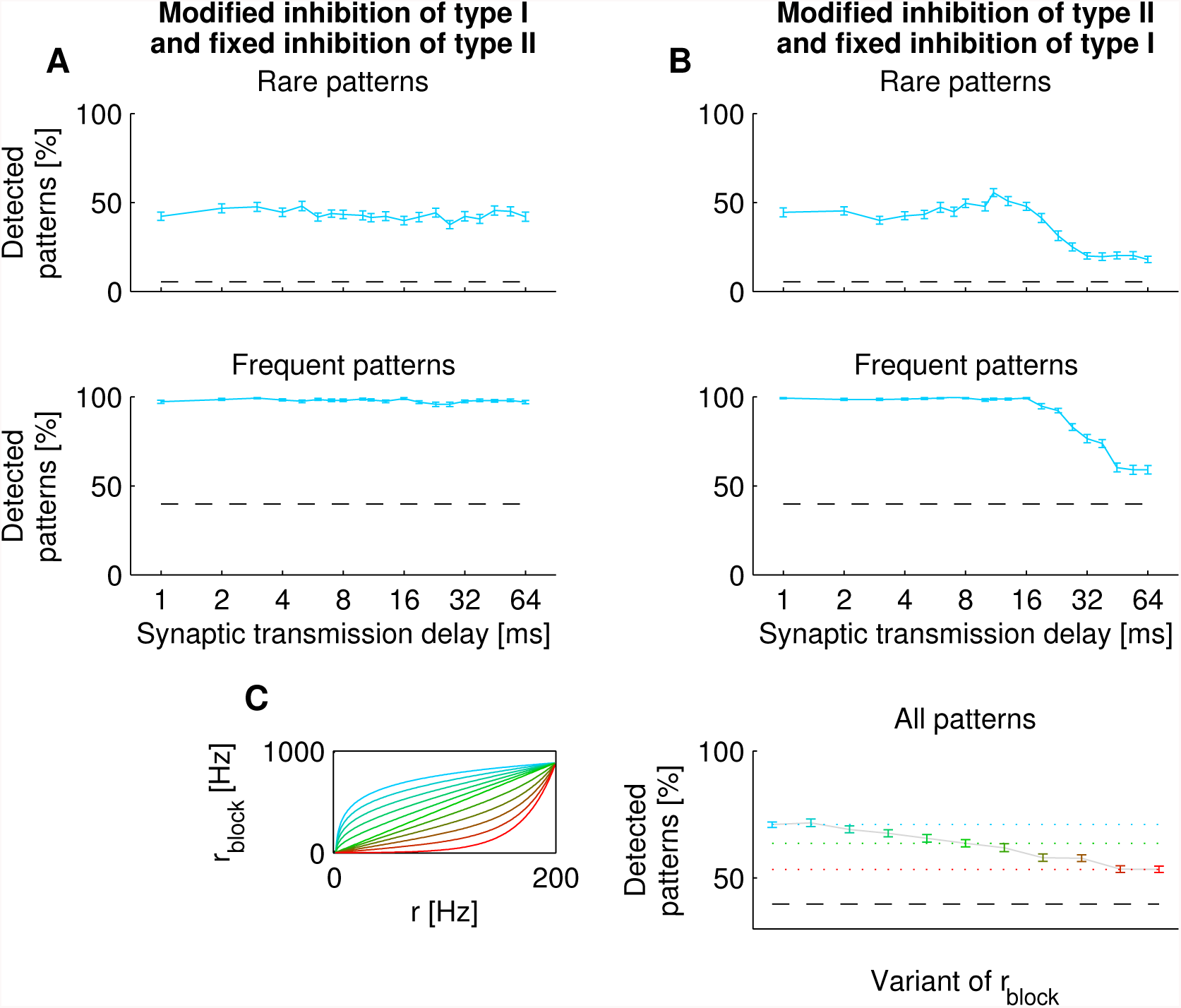
The response of inhibitory neurons neither has to be accurate nor precisely timed. Performance results for WTA networks that consist of twelve PCs and implement inhibition of type I and II. **A:** Dependence of the percentage of detected firing rate patterns on the synaptic transmission delay for inhibition of type I. The synaptic transmission delay for inhibition of type II is set to 1 ms. **B:** Dependence of the percentage of detected firing rate patterns on the synaptic transmission delay for inhibition of type II. The synaptic transmission delay for inhibition of type I is set to 1 ms. Only for a synaptic transmission delay of more than 16 ms the performance drops. The performance is not affected by imprecise synaptic transmission delays. **C:** Right panel: Dependence of the performance on the shape of the total firing rate function of all inhibitory neurons of type II *r*_block_. The results for different variants of *r*_block_ are indicated with the color of the error bars. The color code is defined in the left panel. Left panel: Different variants of *r*_block_ in dependence of the average firing rate of all PCs *r* are shown in different colors. The theoretically optimal variant is indicated in light blue. All results are shown for WTA networks that implement medium strength feedback inhibition of type I (*r*_inh_ = 2*r*). Dashed lines indicate the performance results for WTA networks without inhibition of type II and a synaptic transmission delay of 1 ms (see Fig. 4). Inhibitory neurons of type II with a linear firing rate function *r*_block_ improve the performance by 25% when compared to WTA networks without inhibition of type II.

For inhibitory neurons that provide inhibition of type II only a synaptic transmission delay larger than 16 ms causes a performance drop (Fig. 5B). Furthermore, small distortions of the firing rate function of neurons that provide inhibition of type II have only a small impact on the performance (Fig. 5E). It is sufficient that the firing rate function is approximately a logarithmic function of the total firing rate of PCs. A linear firing rate function still improves the performance of the WTA network by 25% compared to a WTA network without inhibition of type II. Finally, the performance does not depend on the number of synaptic connections between PCs and inhibitory neurons, as long as the connection probability stays above 50% (Fig. 6). This holds true for WTA networks that implement inhibition of type I and WTA networks that implement both types of inhibition. In summary, inhibition neither has to be accurate nor precisely timed to result in high performance values.

**Figure 6:**
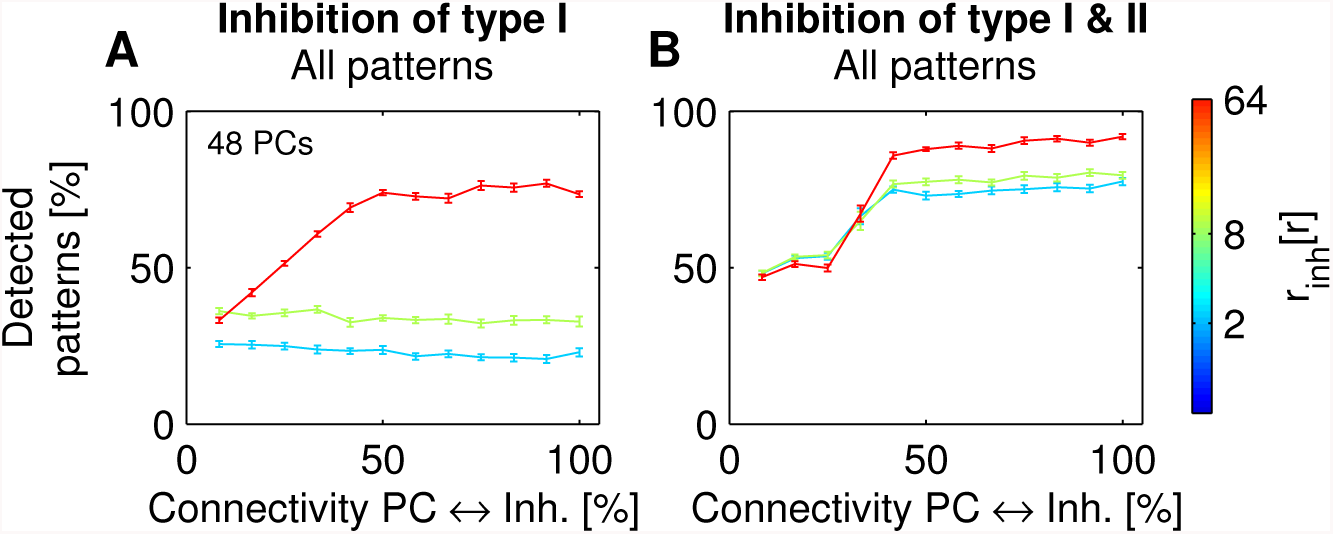
Performance in dependence on the connection probability between PCs and inhibitory neurons. Dependence of the performance on the percentage of established synaptic connections between 48 PCs and 12 inhibitory neurons for WTA networks the implement inhibition of type I (**A**), and inhibition of type I and II (**B**). The average total firing rate of all inhibitory neurons of type I *r*_inh_ is set to a multiple of the average firing rate of all PCs *r*. The synaptic weights are rescaled so that average synaptic excitatory and inhibitory inputs stay constant. The performance drops for connection probabilities below 50%.

### 3.5 Lateral synaptic connections induce experimentally observed neural correlations

Superficial cortical layers are considered to be the primary candidate for a possible implementation of WTA networks (Douglas and Martin, 2004). Experimental data on spontaneous and sensory-evoked neocortical activity in-vivo has shown that PCs in superficial cortical layers discharge correlated in a spatially organized manner (Sakata and Harris, 2009). Correlations between the activity of local pairs of PCs (separated by < 50*μm*) are much stronger than for distal pairs (separated by > 200*μm*). These results seem to be in conflict with the sparse activity of PCs in WTA networks. We investigate whether the experimentally observed correlations might be attributed to local lateral excitatory synaptic connections between PCs, which are not included in conventional WTA networks.

In order to induce the observed correlations between the activity of local pairs of PCs we include local lateral synaptic connections in a WTA network by pooling PCs to local patches consisting (of disjoint subsets) of four neurons. PCs within the same patch are fully connected with uniform synaptic strength. Pairs of PCs within the same patch and across patches are considered to be local and distal, respectively. We carried out a correlation analysis on spike counts in the 50 ms period following firing rate pattern onsets as in (Sakata and Harris, 2009). We analyzed correlations between average responses of PCs to different firing rate patterns, called *signal correlations.* In addition, we investigated correlations between responses to repetitions of the same firing rate pattern, called *noise correlations.* Both types of correlations are larger for local pairs of PCs than for distal pairs (Fig. 7). Moreover, for local pairs both types of correlations can be set to arbitrary values by adjusting the strength of the lateral synaptic connections (Fig. 7C). In contrast, the corresponding values for distal pairs of PCs remain close to zero (with correlation coefficient smaller than 0.1). Overall, the results for WTA networks that implement lateral excitatory synaptic connections are consistent with experimental data on neocortical activity.

**Figure 7:**
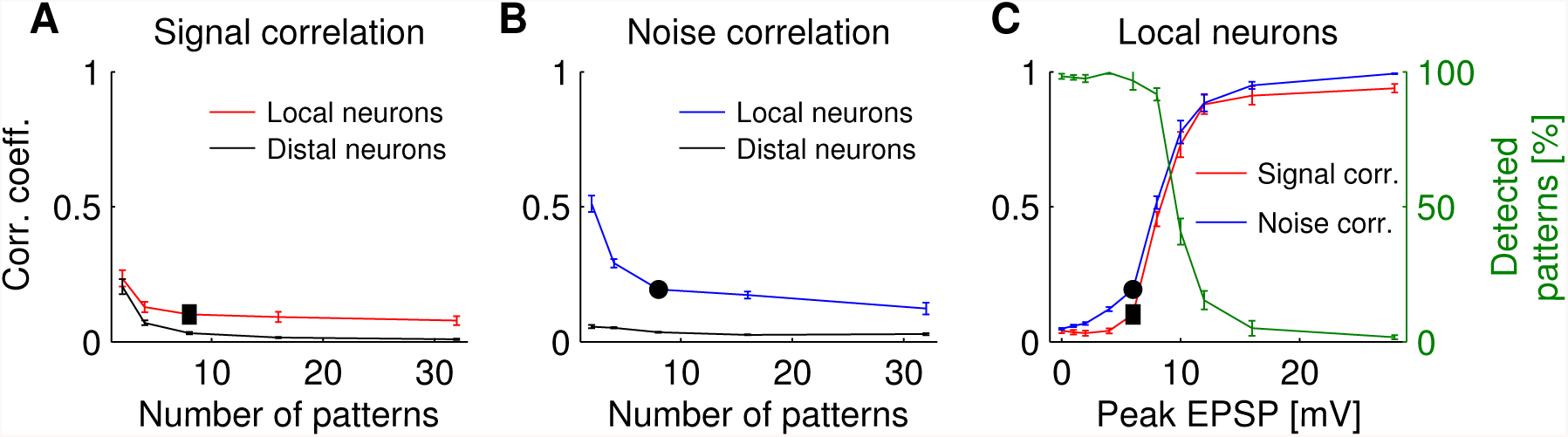
Spatial dependence of spike count correlations. Correlation analysis on spike counts in the 50 ms period following firing rate pattern onset. Results are shown for WTA networks consisting of 48 PCs that implement medium strength inhibition of type I (r_in_h = 2r) and inhibition of type II for various numbers of firing rate patterns. Local lateral synaptic connections are implemented by pooling PCs to local patches consisting (of disjoint subsets) of four neurons. Signal correlations (**A**) and noise correlations (**B**) are larger for local pairs (within the same patch) than for distal pairs (across patches). For local pairs both correlations can be set to arbitrary values by adjusting the strength of lateral synaptic connections (**C**). Only for large EPSPs (> 6 mV) the percentage of detected patterns drops. The results are consistent with experimental data on correlations of neural firing during spontaneous and sensory-evoked neocortical activity (Sakata and Harris, 2009).

## 4 Discussion

The contribution of different anatomical subtypes of interneurons in cortical microcircuits (reviewed in Markram et al., 2004; Ascoli et al., 2008) to the emergence of cortical function remains unclear. Our analysis suggests a new theoretical perspective on the functional role of dendritic-targeting interneurons: They implement competition in ensembles of PCs by normalizing the rate of synaptic plasticity. Previous theoretical work has shown that STDP in combination with symbolic lateral inhibition installs in ensembles of PCs probabilistic models of their input. But these models rely on a normalized firing rate of PCs that is difficult to achieve with spiking inhibitory interneurons. We have shown that this normalization of the activity of PCs can be replaced by a normalization of the rate of STDP. We propose a model with an inhibitory network that consists of two types of interneurons that are distinguished by the structure of their axonal arborization, i.e. they target either somatic or dendritic compartments of PCs.

The most frequent dendritic-targeting interneurons are somatostatin expressing Martinotti cells. Although it is generally accepted that synaptic input from Martinotti cells influences dendritic processing and synaptic plasticity, it has been difficult to identify their precise function. We hypothesize that strong synaptic input from Martinotti cells blocks STDP in the distal dendrites of PCs such that that the expected synaptic weight change of post-synaptically triggered STDP is independent of the total firing rate of the ensemble of PCs. We conjecture that the blocking of STDP is implemented by intercepting backpropagating action potentials. It is well known that the induction of long-term potentiation (LTP) depends on the co-activation of excitatory post synaptic potentials and back-propagating action potentials (Yuste and Denk, 1995; Magee and Johnston, 1997; Koester and Sakmann, 1998; Schiller et al., 1998). If backpropagating action potentials fail, NMDA receptors might not be sufficiently depolarized to provide enough calcium influx to trigger LTP (Artola et al., 1990; Sjostrom and Hausser, 2006). In addition, Bar-Ilan et al. (2012) have shown that excitatory synaptic plasticity depends critically on the exact localization and strength of inhibitory synapses. Sufficient strong inhibition in dendrites protects nearby excitatory synapses from undergoing plasticity by suppressing the level of calcium. These observations fit well with the required mechanism for blocking STDP.

The predominant somatic-targeting interneurons are parvalbumin expressing basket cells that presumably normalize the activity of PCs and provide feedback inhibition to balance excitatory synaptic inputs (reviewed by Isaacson and Scanziani, 2011). Okun and Lampl (2008) have shown that excitatory and inhibitory synaptic inputs to pairs of nearby neurons are continuously correlated in strength and inhibition lags behind excitation by about 4 ms. These experimental observations agree well with the results for our model where feedback inhibition lags behind excitation by about 5 ms. Moreover, recent studies in mouse visual cortex *in vivo* (Kwan and Dan, 2012) showed that the two major types of inhibitory networks carry out distinct operations, i.e. somatostatin expressing interneurons detect single-neuron firing and parvalbumin expressing interneurons respond to distributed network activity. These results agree very well with the model assumptions that parvalbumin expressing interneurons carry out a rough normalization of the network activity and somatostatin expressing interneurons implement precise competition between sparsely firing PCs.

A key criterion for the biological plausibility of the WTA network is its agreement with experimental findings on the functional connectivity between PCs and inhibitory neurons. It might seem questionable, whether the two implemented interneuron types provide sufficient strong inhibition for PCs. An experimental study (Kwan and Dan, 2012) estimated that up to 9 somatostatin expressing interneurons are recruited by a single PC that spikes with a firing rate of about 60 Hz in response to a 100 ms stimulus (as in our simulations). The actual number of activated somatostatin expressing interneurons is assumed to be even larger, because of a stringent statistical criterion used to test for functional connectivity. In addition, this work reports a firing rate function for somatostatin expressing interneurons that is consistent with a linear function (see Fig. 5C) with a slope of about 0.3. Based on this estimate and our results for inhibition of type II about 14 Martinotti cells have to respond to spikes of single PCs for successful self-organization. This fits well the experimental data in (Kwan and Dan, 2012). In addition, the model doesn’t require that basket cells are recruited by the firing of single PCs. However, the stronger the modulation of feedback inhibition provided by basket cells the better the performance.

The local anatomical connectivity between inhibitory neurons and PCs in neocortex is still a matter of debate. Several experimental studies based on double or triple patchclamp recordings report probabilities of connections between PCs and somatostatin-expressing interneurons ranging from 20% in layer 2/3 (Thomson and Lamy, 2007; Thomson and Morris, 2002; Thomson et al., 2002; Yoshimura and Callaway, 2005) to 3% in layer 5 (Otsuka and Kawaguchi, 2009). In contrast, Fino and Yuste (2011) estimated probabilities of connections between somatostatin-expressing interneurons and PCs in layer 2/3 with values above 70% based on two-photon photostimulation. We have demonstrated in simulation that connections probabilities above 50% are sufficient to reliably induce probabilistic inference in ensembles of PCs.

A testable prediction of the model is that silencing either Martinotti cells or basket cells reduces but does not abolish the emergence of sparse codes in ensembles of PCs. Moreover, if the firing patterns of upstream neurons can be recorded (or evoked) and the firing rates of all active PCs can be increased by a fixed amount during the representations of a single pattern, then silencing either Martinotti cells or basket cells has a different effect on the sparseness of the neural code (after learning). Active Martinotti cells and silent basket cells result in the same fraction of active PCs during the presentation of each pattern. In contrast, active basket cells and silent Martinotti cells result in an increased number of active PCs. In conclusion, we have demonstrated how two inhibitory networks can orchestrate the self-organization of computational function in cortical microcircuits and, thereby, provided new concepts and methods for integrating diverse results from interneurons into a principled framework.

## Acknowledgements

This project/research received funding from the European Unions Horizon 2020 Framework Programme for Research and Innovation under the Framework Partnership Agreement No. 650003 (HBP FPA).

